# Monoclonal antibodies for detecting collagen IV α3α4α5 in basement membranes

**DOI:** 10.64898/2026.11.18.767278

**Authors:** Maryline Fresquet, Gema Bolas, Emily Williams, Shota Kaseda, Holly Birchenough, Sherylanne Newton, Conor J. Sugden, Thomas A. Jowitt, Andrew Steinsapir, Kevin A. Goncalves, Daniel J. Jagger, David R. Sherwood, Rachel Lennon

## Abstract

The collagen IV α3α4α5 trimer forms a specialised network within basement membranes that is essential for normal kidney, inner ear and ocular function. Variants in *COL4A3*, *COL4A4* or *COL4A5* disrupt this network and cause Alport syndrome. Yet, current antibody tools detect individual collagen IV chains rather than the assembled α3α4α5 trimer. Here, we generated a recombinant human mini-COL4A345 trimer and used it to produce mouse monoclonal antibodies that recognise native collagen IV α3α4α5.

We screened hybridomas using biolayer interferometry, native dot blotting, denaturing western blotting and immunofluorescence, followed by characterisation of selected purified antibodies using surface plasmon resonance. Three clones, Ms#5, Ms#44 and Ms#72, detected native, folded collagen IV α3α4α5. We validated antibody specificity using *Col4a3*, *Col4a5* and *Col4a6* knockout mouse tissues, and assessed their utility for human kidney immunofluorescence and immunoprecipitation from mouse kidney.

The antibodies detected conformationally intact collagen IV α3α4α5 in tissue, with no detectable cross-reactivity with collagen IV α1α1α2 or α5α5α6 networks under the conditions tested. Clone Ms#72 provided the strongest and most consistent tissue signal, localising collagen IV α3α4α5 to the glomerular basement membrane, cochlear basement membranes and retinal pigment epithelium-associated basement membrane. Ms#72 also immunoprecipitated endogenous collagen IV α3, α4 and α5 from wild-type kidney tissue, confirming recognition of the native complex.

These monoclonal antibodies provide a resource for detecting collagen IV α3α4α5 in basement membrane networks. Their potential applications include assessment of the α3α4α5 heterotrimer in patient biopsies and evaluation of gene therapies and other interventions aimed at modulating α3α4α5 deposition in Alport syndrome.

## Introduction

Basement membranes are specialised extracellular matrices that surround most tissues, provide structural support and regulate cell behaviour. Type IV collagen is composed of six genetically distinct α chains (α1- α6), which assemble into at least three trimeric protomers: α1α1α2, α3α4α5 and α5α5α6. These protomers assemble into networks within basement membranes [1, 2]. The α3α4α5 network is a tissue-restricted collagen IV network required for specialised basement membrane function. It is encoded by *COL4A3*, *COL4A4* and *COL4A5*, and pathogenic variants in these genes impair network formation and cause Alport syndrome, a disorder characterised by progressive kidney disease, hearing loss and ocular abnormalities [3–6]. The three α chains share a conserved domain organisation, with an amino-terminal signal peptide, a 7S domain, a long collagenous Gly-X-Y repeat region and a carboxy-terminal non-collagenous NC1 domain [7, 8]. Type IV collagen α chains are highly conserved between humans and mice, supporting the use of mouse models to evaluate collagen IV network assembly, tissue localisation and function.

Despite its biological and clinical importance, the assembled collagen IV α3α4α5 network has been difficult to measure directly. Available antibodies are raised against short peptide sequences from the NC1 domains of individual collagen IV chains and therefore report chain presence rather than heterotrimer assembly. Consequently, network formation is often inferred from co-localisation of α3, α4 and α5 chain-specific immunodetection [9]. This approach cannot distinguish whether the α3, α4 and α5 chains have assembled into a mature heterotrimeric network, whether individual chains have assembled correctly, or whether therapeutic strategies have restored the α3α4α5 network.

Direct detection of assembled α3α4α5 would provide a readout of basement membrane maturation that is not captured by chain-specific labelling alone. A monoclonal antibody recognising a conformational epitope within the native folded α3α4α5 heterotrimer could distinguish loss of heterotrimer assembly from reduced expression or mislocalisation of individual α3, α4 or α5 chains. It could also quantify residual or restored α3α4α5 deposition in kidney tissue, test whether therapeutic delivery of one or more collagen IV chains reconstitutes the endogenous network, and provide a translational assay for evaluating gene replacement, exon-skipping, RNA-based or protein-engineering approaches designed to rebuild the α3α4α5 network in basement membranes.

Here, we describe the development and validation of mouse monoclonal antibodies raised against a recombinant human mini-COL4A345 antigen. We prioritised antibodies that recognise the folded trimer, then tested candidate clones using biochemical binding assays, human and mouse tissue immunofluorescence, knockout mouse controls and immunoprecipitation from solubilised glomeruli. This work provides a renewable antibody resource for assessing collagen IV α3α4α5 in basement membranes.

## Results

### Development of a mini-COL4A345 trimer antigen

Recombinant production of full-length collagen IV α3α4α5 is technically challenging; therefore, we designed a mini-COL4A345 antigen containing the three constituent collagen IV α3, α4 and α5 chains (**Supplementary Figure 1**). Each construct retained the NC1 domain, which drives chain selection and trimer assembly, and the 7S domain, which contributes to collagen IV network formation (**Figure 1A**). We expressed and purified the recombinant protein to generate a mini-COL4A345 preparation suitable for biochemical characterisation. Mass spectrometry confirmed protein identity and purity, and peptide mapping defined the sequence coverage. SDS-PAGE resolved bands of approximately 40 kDa under reducing conditions, corresponding to the individual mini-COL4A3, -COL4A4 and -COL4A5 chains. Under non-reducing conditions, the predominant species migrated at approximately 110 kDa, consistent with a mini-COL4A345 heterotrimer, with a smaller higher-molecular-weight species at approximately 240 kDa, consistent with hexamer formation (**Figure 1B**). Sedimentation velocity analytical ultracentrifugation identified mini- COL4A345 as predominantly trimeric in solution, with an estimated molecular mass of approximately 126 kDa and only a minor higher-order population consistent with hexamers (**Figure 1C**). Mass photometry independently supported this oligomeric distribution (**Supplementary Figure 2**). Finally, an AlphaFold-predicted model recapitulated the expected trimeric organisation of the mini-COL4A345 complex (**Figure 1D**). Together, these data confirm that the recombinant mini-COL4A345 antigen predominantly assembles as a trimer and was therefore suitable for generating monoclonal antibodies against the heterotrimer.

**Figure 1.**
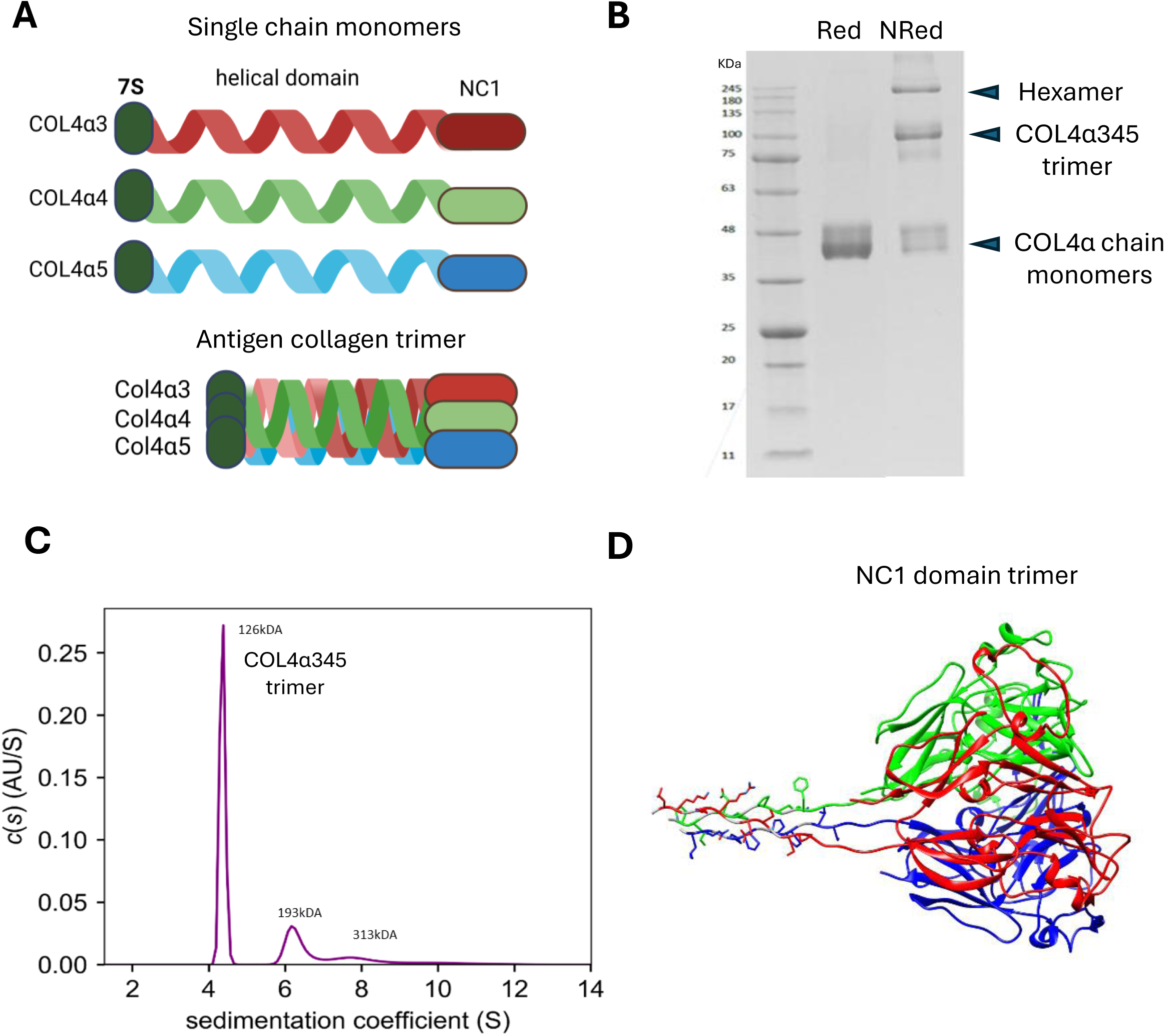
Characterisation of the mini-COL4A345 trimer antigen. A) Schematic representation of collagen IV α3α4α5 heterotrimer assembly. Individual COL4A3, COL4A4 and COL4A5 chains contain an N-terminal 7S domain, a triple-helical collagenous domain and a C-terminal NC1 domain. The mini-COL4A345 antigen contains all three chains with a shortened collagenous region and assembles through the NC1 domains. B) Coomassie-stained SDS-PAGE gel of purified mini-COL4A345. Under reducing conditions, COL4 monomers migrated at approximately 40 kDa. Under non-reducing conditions, the complex migrated predominantly at approximately 110 kDa, consistent with a trimer, with a higher molecular weight species at approximately 240 kDa, consistent with a hexamer. Red, reduced; NRed, non-reduced. C) Sedimentation velocity analytical ultracentrifugation of purified mini-COL4A345 confirming the predominant trimeric species in solution. D) AlphaFold-predicted structure of the mini-COL4A345 antigen. Mini-COL4A3, COL4A4 and COL4A5 are shown in red, green and blue, respectively, and fold into the expected NC1 trimer.

### Generation and characterisation of COL4A345 monoclonal antibodies

We immunised mice with the purified mini-COL4A345 antigen, generated hybridomas and screened them against the recombinant proteins. We assessed the 40 highest-performing parental clones from the initial ELISA screen for binding to purified mini-COL4A345, mini- COL4A3 and mini-COL4A5 using biolayer interferometry (BLI) and immunoassays. Fourteen clones bound the mini-COL4A345 trimer, with responses classified as “high” or “low” binders based on binding response and signal above reference-subtracted baseline (**Supplementary Figure 3A**). Under denaturing western blotting conditions, three hybridomas (clones #21, #37 and #85) reacted with mini-COL4A345 and were excluded from further selection, as we sought antibodies that preferentially recognised the assembled COL4A345 trimer rather than denatured COL4 chains. Two hybridomas (clones #72 and #79) recognised native collagen IV α3α4α5 by immunofluorescence in human kidney tissue (**Supplementary Figure 3B**). Of the remaining 26 clones with no detectable mini-COL4A345 binding by BLI, one recognised denatured mini-COL4A345 (clone #43) while three detected native collagen IV α3α4α5 by immunofluorescence (clones #5, #44 and #70). Clones #5, #44 and #72 showed strong glomerular basement membrane localisation and were selected for monoclonal antibody production (**Supplementary Figure 3B**). We characterised purified monoclonal antibodies Ms#5, Ms#44 and Ms#72 by surface plasmon resonance (SPR). BLI enabled rapid initial screening of hybridomas, while SPR was used for subsequent kinetic characterisation, allowing more detailed assessment of association and dissociation kinetics and binding affinity. Kinetic measurements against immobilised recombinant mini-COL4A345 and the individual recombinant mini-COL4A3 and -COL4A5 chains showed that Ms#5 and Ms#44 selectively recognised the mini-COL4A345 heterotrimer, with no detectable binding to individual mini-COL4A3 or -COL4A5 chains. Recombinant mini-COL4A4 could not be generated and therefore was not tested. Ms#72 bound strongly to mini-COL4A345 and also to mini-COL4A3 (**Figure 2A,B**). Together, these results identified three monoclonal antibodies that recognise folded mini-COL4A345, with Ms#72 showing the strongest overall performance across biochemical and tissue-based immunofluorescence assays.

**Figure 2.**
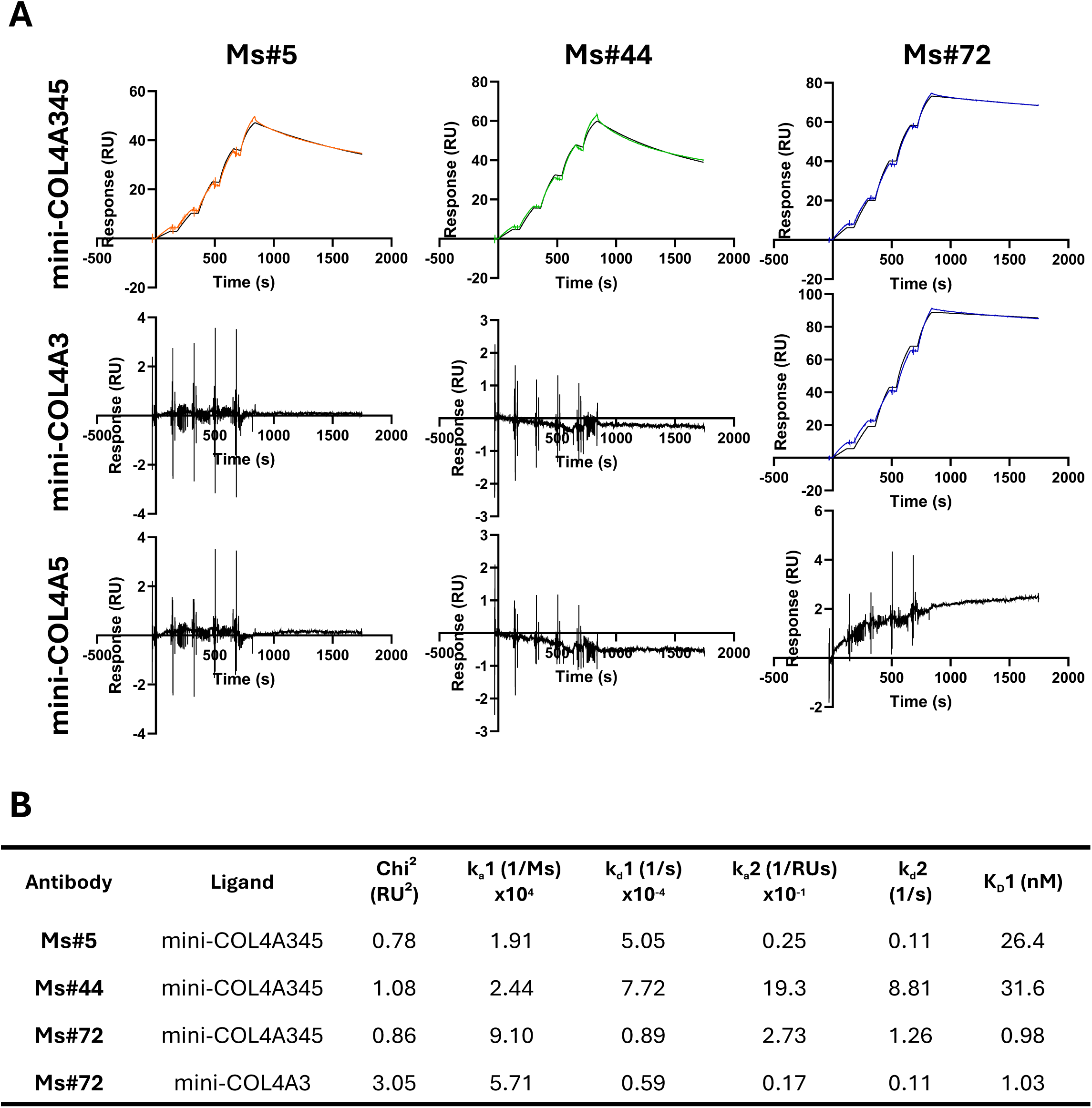
Binding analysis of selected monoclonal antibodies to mini-COL4A345, - COL4A3 and -COL4A5. A) Surface plasmon resonance sensorgrams for binding of Ms#5, Ms#44 and Ms#72 to immobilised recombinant mini-COL4A345, -COL4A3 and -COL4A5. Antibody binding response shown in coloured lines and bivalent analyte model fit shown in black. B) Summary of calculated kinetic parameters from SPR analysis.

### Assessment of antibody specificity using knockout mouse models

We next assessed antibody specificity by immunofluorescence in kidney tissue from wild- type mice and from *Col4a3*, *Col4a5* and *Col4a6* knockout models. Ms#5, Ms#44 and Ms#72 labelled the glomerular basement membrane and tubular basement membrane strongly in wild-type FFPE sections (**Figure 3A**; **Supplementary Figure 4A**) and kidney cryosections (**Figure 3B**; **Supplementary Figure 4B**). In contrast, *Col4a3* and *Col4a5* knockout kidneys lacked an immunofluorescence signal, demonstrating that antibody binding depends on the presence of the collagen IV α3α4α5 heterotrimer (**Figure 3A,B**). In *Col4a6* knockout kidneys, the antibodies labelled the glomerular basement membrane but not Bowman’s capsule, indicating that antibody recognition is independent of the α6 chain and does not detect the α5α5α6 network (**Figure 3A**). These data show that the monoclonal antibodies specifically recognise collagen IV α3α4α5 in tissue sections.

**Figure 3.**
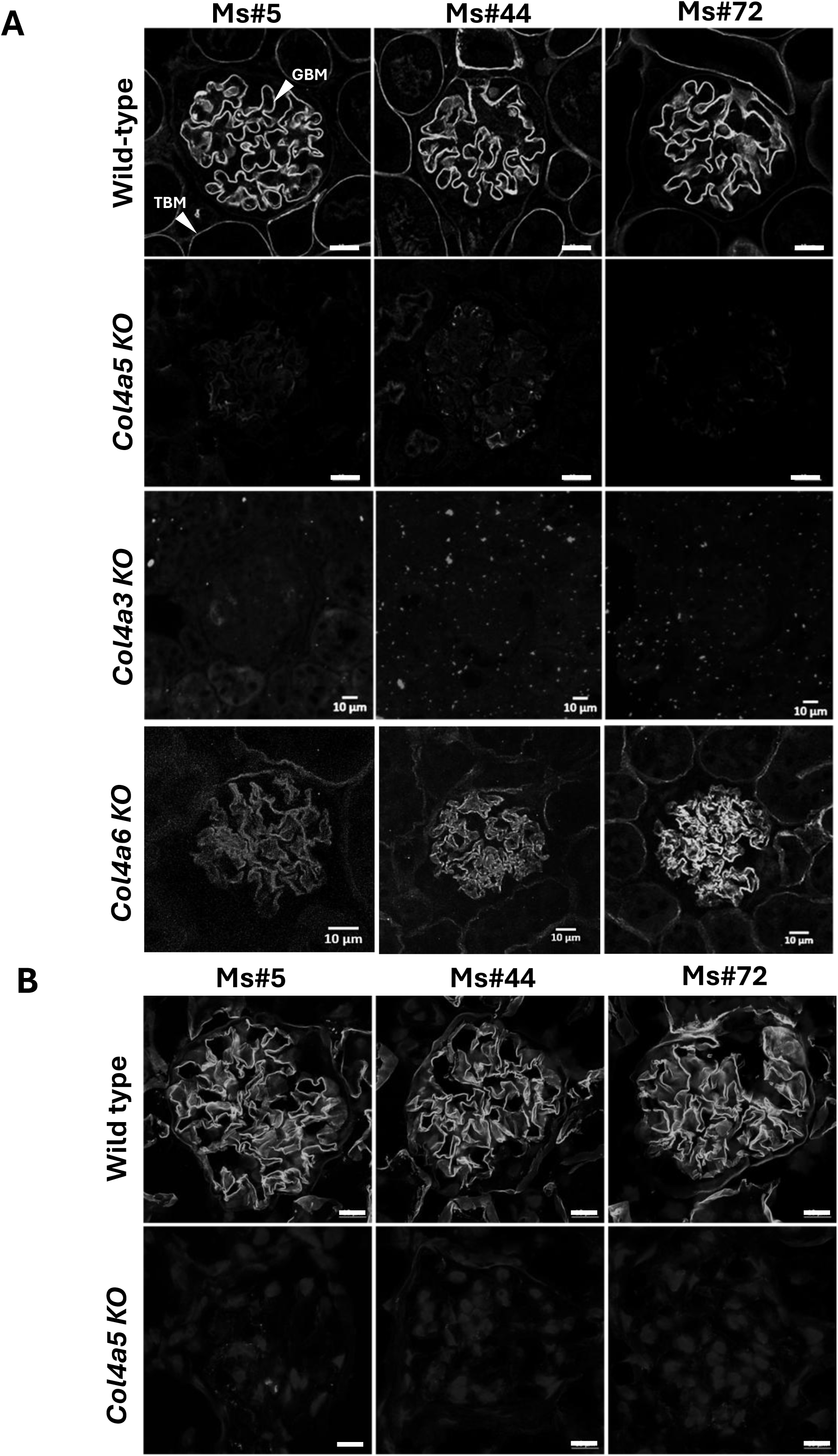
Evaluation of antibody specificity using knockout mouse models. A) FFPE mouse kidney sections stained with monoclonal antibodies Ms#5, Ms#44 and Ms#72. Antibodies labelled the glomerular basement membrane (GBM) and tubular basement membrane (TBM) in wild-type and *Col4a6* knockout mice, whereas immunofluorescence was absent from *Col4a3* and *Col4a5* knockout tissues. B) Mouse kidney cryosections stained with Ms#5, Ms#44 and Ms#72 showed the same collagen IV α3, α4 and α5-dependent GBM and TBM localisation pattern. Scale bars, 10 μm.

### Specific localisation of anti-COL4A345 monoclonal antibodies in human kidney

We then evaluated antibody performance in human kidney tissue. In healthy human kidney, Ms#5, Ms#44 and Ms#72 labelled the glomerular basement membrane, consistent with collagen IV α3α4α5 localisation (**Figure 4A**). Co-labelling with chain-specific antibodies demonstrated that Ms#72 co-localised with the collagen IV α3, α4 and α5 chains in the glomerular basement membrane (**Figure 4B**). In contrast, Ms#72 did not co-localise with the collagen IV α1 or α2 chains, supporting selective recognition of the α3α4α5 trimer rather than the ubiquitous α1α1α2 network (**Supplementary Figure 5**). Of note, Ms#72 also showed weak partial labelling of Bowman’s capsule under these conditions, in agreement with recent findings demonstrating the presence of collagen IV α3 and α5 in Bowman’s capsule [10]. To determine whether the Bowman’s capsule signal reflected cross-reactivity with collagen IV α5α5α6, FFPE human kidney sections were co-labelled with Ms#72 and antibody against the collagen IV α6 chain. Collagen IV α6 immunofluorescence was strong in Bowman’s capsule but absent from the glomerular basement membrane (**Figure 4B**).

**Figure 4.**
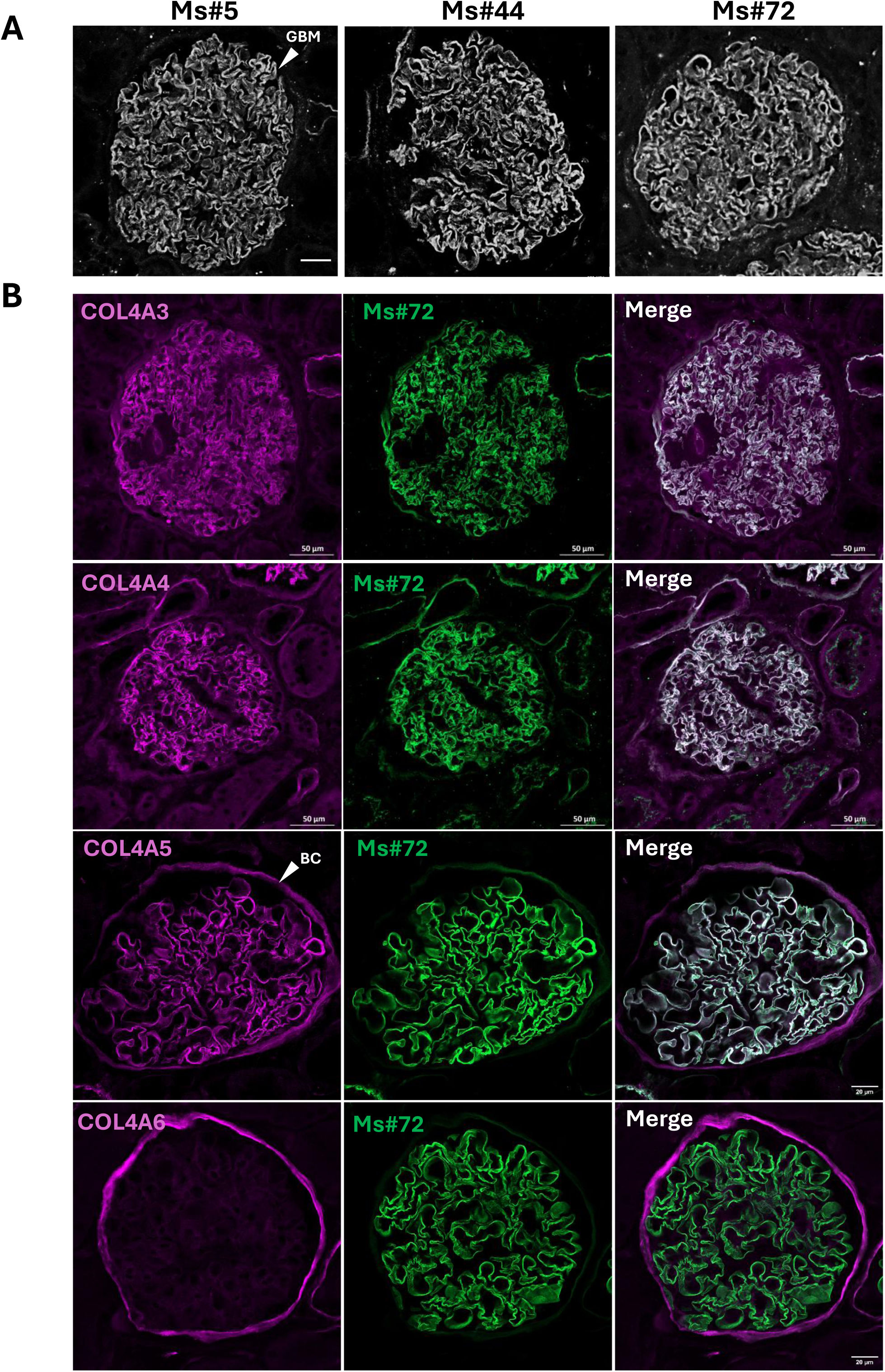
Validation of antibody specificity in human kidney by immunofluorescence. A) Representative images of FFPE human kidney sections stained with Ms#5, Ms#44 and Ms#72, showing native collagen IV α3, α4 and α5 localisation in the glomerular basement membrane (GBM). Scale bar, 50µm. B) Human glomerular sections co-labelled with Ms#72 and chain-specific antibodies against COL4A3, COL4A4, COL4A5 or COL4A6. Ms#72 co- localised strongly with COL4A3, COL4A4 and COL4A5 in the GBM, whereas minimal overlap was observed with COL4A6. BC, Bowman’s capsule. Scale bars, 50 μm (top panels) and 20 μm (bottom panels).

Overall, these data support detection of collagen IV α3α4α5 in the human glomerular basement membrane without detectable cross-reactivity with α5α5α6 in Bowman’s capsule.

### Localisation of COL4A345 in cochlea and eye

Since Ms#72 had the most consistent signal in tissue, we performed further immunofluorescence analysis with this antibody alone. We tested whether Ms#72 could detect collagen IV α3α4α5 in the basement membranes of the cochlea and eye, which are also affected in Alport syndrome. In the wild-type cochlea, Ms#72 labelled the lateral wall, including root cells and blood vessels, and the spiral limbus, stria vascularis and basilar membrane (**Figure 5A**, **upper panels**). Signal was absent from inner hair cells, outer hair cells and non-sensory supporting cells within the organ of Corti. This distribution is consistent with previous studies using antibodies against the individual collagen IV α3, α4 and α5 chains [11–13]. The labelling described above was absent in *Col4a5* knockout cochleae (**Figure 5A**, **lower panels**). In the eye, Ms#72 immunofluorescence was restricted to the retinal pigment epithelium adjacent to Bruch’s membrane, with no signal detected in other retinal layers or the sclera (**Figure 5B**). This ocular pattern is consistent with previous reports of collagen IV localisation in Alport models [14]. Together, these findings show that Ms#72 detects collagen IV α3α4α5 in the cochlea and eye in addition to the kidney.

**Figure 5.**
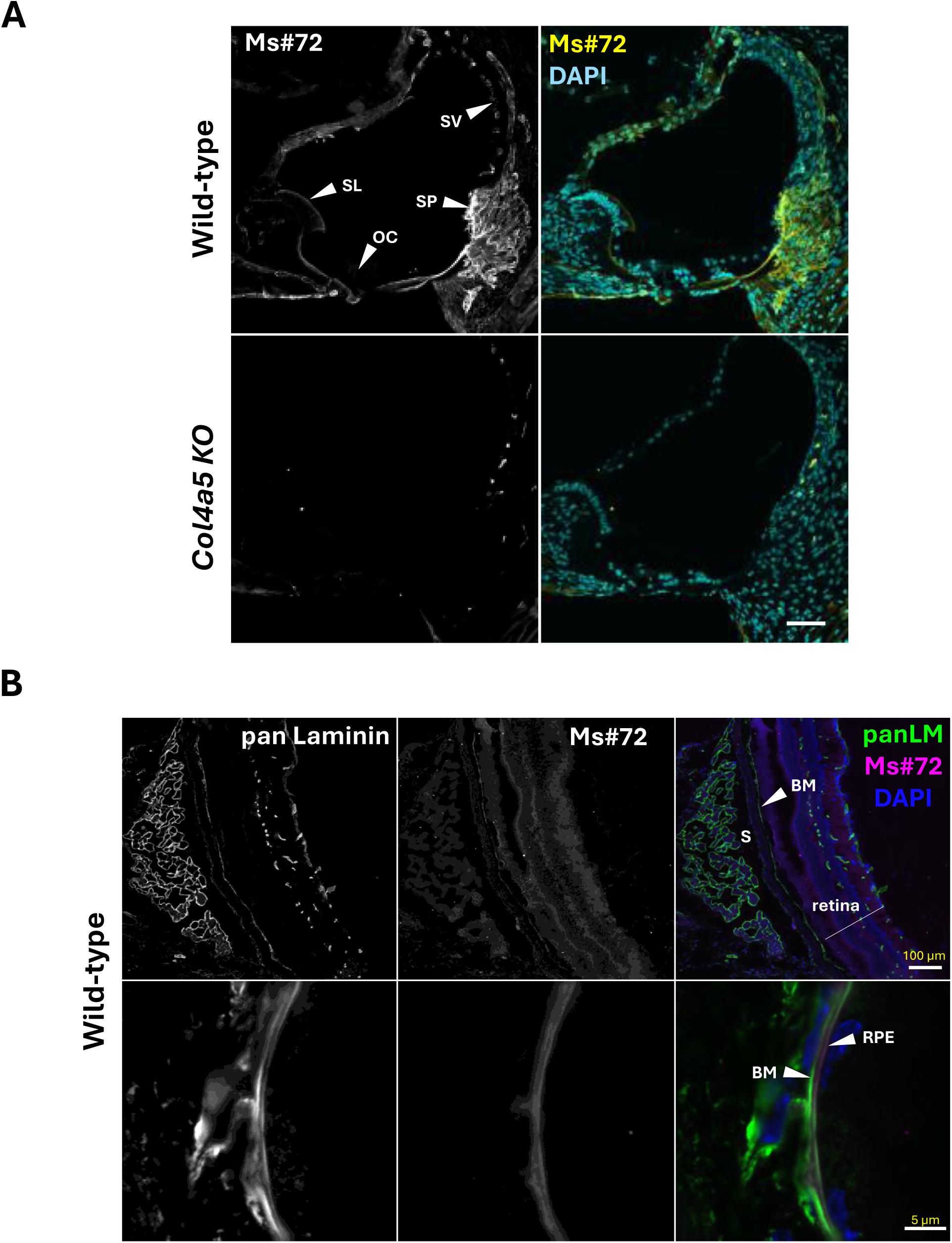
Native collagen IV α3, α4 and α5 localisation in cochlea and eye using Ms#72. A) Cochlear cryosections from 3-week-old wild-type and *Col4a5* knockout mice stained with Ms#72. In wild-type cochlea, COL4A345 labelling was detected in the spiral limbus (SL), stria vascularis (SV), and lateral wall behind the spiral prominence (SP). Staining was absent from the organ of Corti (OC). In age-matched *Col4a5* knockout littermates, Ms#72 signal was absent except for non-specific labelling of blood cells. Scale bar, 50 μm. B) Eye immunofluorescence showed COL4A345 localisation restricted to the retinal pigment epithelium (RPE) adjacent to Bruch’s membrane (BM). No staining was observed in other retinal layers or in the sclera (S). Scale bars, 100 μm (top panel) and 5 μm (bottom panel).

### Validation of Ms#72 binding to native COL4A345

Finally, we tested whether Ms#72 could capture endogenous collagen IV α3α4α5 from kidney tissue. We used collagenase to digest kidney tissue and generate homogenates from wild-type and *Col4a5* knockout mice, then performed immunoprecipitation with Ms#72 (**Figure 6A**). Western blotting analysis of the immunoprecipitated material detected the collagen IV α3, α4 and α5 chains in wild-type samples, demonstrating that Ms#72 captures the native α3α4α5-containing complexes from kidney tissue (**Figure 6B**). *Col4a5* knockout samples lacked α5 and α6, as expected, confirming assay specificity. Detection of higher and lower molecular weight species consistent with previously described monomeric and dimeric collagen IV NC1 fragments [15] further supported the ability of Ms#72 to recover physiologically relevant forms of collagen IV α3α4α5 from mouse kidney tissue.

**Figure 6.**
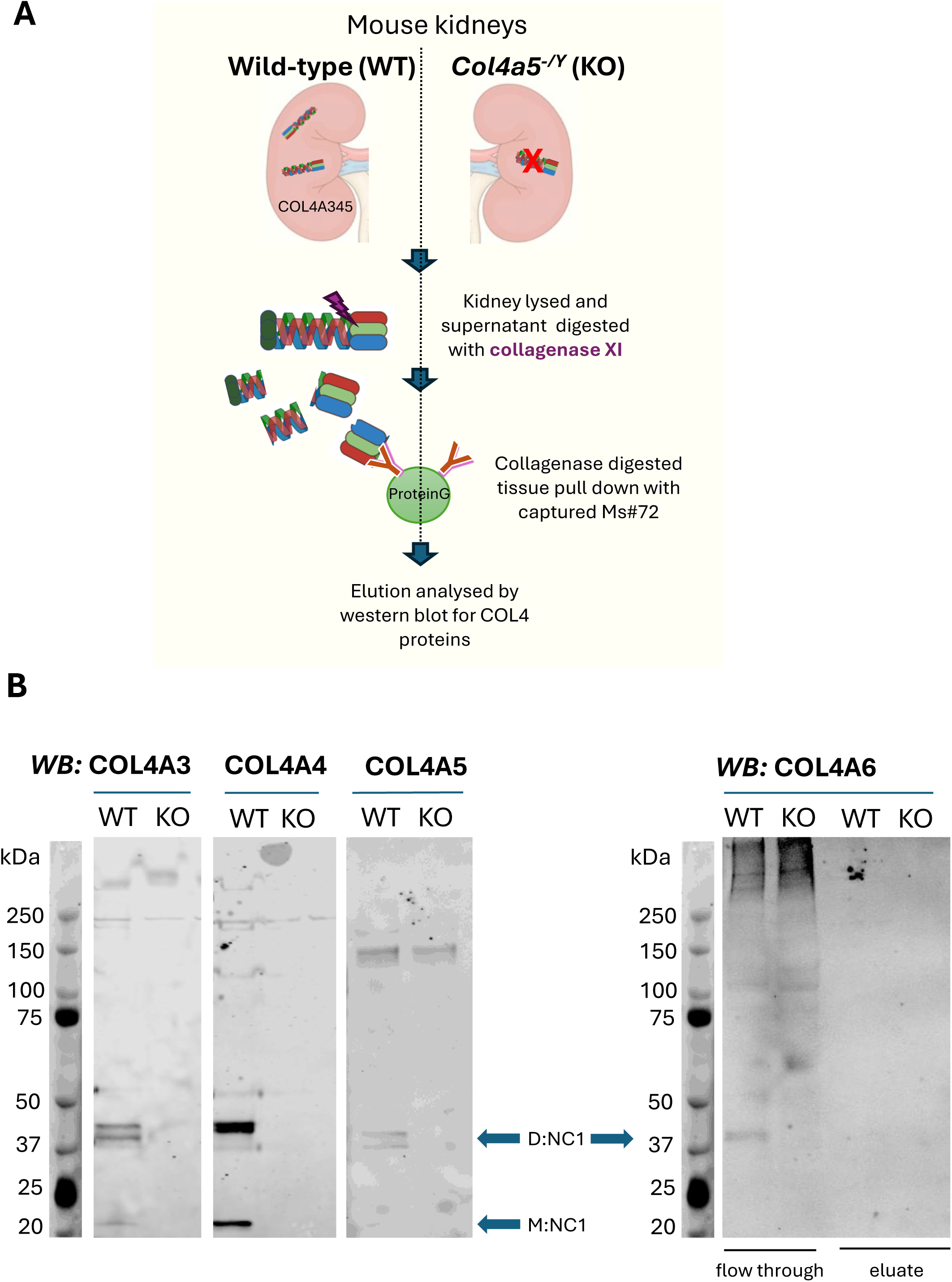
Immunoprecipitation of native collagen IV α3, α4 and α5 from mouse kidney using Ms#72. A) Schematic overview of the immunoprecipitation workflow. Kidney lysates were collagenase-digested and incubated with Ms#72, and antibody-bound protein complexes were captured using Protein G beads before elution and downstream analysis. B) Western blot detection of COL4A3, COL4A4, COL4A5 and COL4A6 in collagenase-digested kidney homogenates from wild-type and *Col4a5* knockout mice after immunoprecipitation with Ms#72. Collagen IV fragments, NC1 dimers referred as D:NC1 and monomers M:NC1 indicated with arrows.

## Discussion

We developed and characterised mouse monoclonal antibodies that detect the collagen IV α3α4α5 trimer in human and mouse tissues. This resource addresses an important gap in the field, as most available reagents detect individual collagen IV chains rather than a conformational epitope present within the native folded α3α4α5 trimer molecule. Monoclonal antibody production provides a renewable and consistent reagent supply, reducing batch-to- batch variability and supporting reproducible assessment of α3α4α5 trimer deposition.

We established antibody specificity using complementary biochemical and tissue-based approaches. *Col4a3* and *Col4a5* knockout kidneys lacked an immunofluorescence signal, demonstrating that antibody binding requires the assembled collagen IV α3α4α5 heterotrimer. *Col4a6* knockout mouse tissue retained a signal in the glomerular basement membrane but lacked a signal in Bowman’s capsule, supporting specificity for collagen IV α3α4α5 rather than the related α5α5α6 heterotrimer. Ms#72 also immunoprecipitated the collagen IV α3, α4 and α5 chains from collagenase-digested kidney tissue, demonstrating recognition of native α3α4α5 complexes. An important next step will be to identify the antibody epitope and define its structural context within the α3α4α5 trimer, potentially through cryo-EM analysis.

Among the antibodies tested, Ms#5 and Ms#44 preferentially recognised the assembled trimeric complex. Ms#72 showed broader reactivity, binding both mini-COL4A345 and the individual mini-COL4A3 chain but demonstrated the strongest and most consistent performance across tissue-based applications. It detected collagen IV α3α4α5 in the glomerular basement membrane and in basement membrane-rich structures in the cochlea and eye, while showing no detectable cross-reactivity with the collagen IV α1, α2 or α6 chains under the conditions tested. Recognition of the assembled trimeric complex provides an advantage over chain-specific immunofluorescence because it gives a direct readout of folded α3α4α5 protomer formation rather than inferred co-localisation of individual chains.

These antibodies have important translational applications. In patient biopsies, they could help determine whether the α3α4α5 heterotrimer is absent, reduced, mislocalised or restored within basement membranes. They also provide a practical assay for therapeutic studies that aim to increase de novo α3α4α5 deposition, including gene replacement, RNA- based, exon-skipping and protein-engineering strategies. In this context, direct detection of the assembled trimer will be more informative than detection of individual collagen IV α3, α4 or α5 chains alone.

The ability of Ms#72 to detect collagen IV α3α4α5 outside the kidney further broadens the use of this resource. Its pattern of immunofluorescence in the cochlea and eye supports applications in studies of hearing and ocular manifestations of Alport syndrome, as well as wider investigations of collagen IV trimer organisation in specialised basement membranes. Together, these monoclonal antibodies provide a defined and renewable toolkit for studying collagen IV biology, evaluating α3α4α5 protomer assembly and supporting preclinical and clinical development of therapies that aim to restore collagen IV α3α4α5 assembly and the network it forms.

In summary, we report a monoclonal antibody resource that enables detection of the folded collagen IV α3α4α5 protomer across relevant tissues. The validation strategy demonstrates specificity for collagen IV α3α4α5 over the related α1α1α2 and α5α5α6 protomers in tissue, compatibility with tissue immunofluorescence and utility for biochemical capture of native α3α4α5. These antibodies could support mechanistic studies of collagen IV assembly, assessment of patient biopsies and future evaluation of therapies designed to increase or restore α3α4α5 deposition.

## Methods

### Expression and purification of COL4 proteins

Mouse monoclonal antibodies were raised against a recombinant human mini-COL4A345 protomer comprising human mini-COL4A3, mini-COL4A4 and mini-COL4A5 proteins (UniProtKB: Q01955, P53420 and P29400). Each modified mini-COL4 protein contained the full native 7S and NC1 domains and a shortened collagenous domain.

Each target insert (mini-COL4A3, -COL4A4 and -COL4A5) was paired with an IgK leader sequence for secretion, a tag for detection or purification (FLAG, TwinStrep or 8x, His), and a TEV protease cleavage site. Expression was driven by the mammalian CMV promoter.

Protein expression and purification were performed by Peak Proteins (Sygnature Discovery, UK). Briefly, IgK-FLAG-TEV-COL4A3, IgK-TwinStrep-TEV-COL4A4 and IgK-8His-TEV-COL4A5 plasmids (**Supplementary Figure 1**) were transiently transfected into HEK293-6E cells (Cellosaurus, CVCL_HF20). Media containing secreted protein were sequentially purified using a 5 mL HisTrap Excel Ni-IMAC column, Strep-Tactin XT affinity chromatography and a HiPrep 26/10 Desalting column.

### Sedimentation velocity analytical ultracentrifugation

The hydrodynamic properties and oligomeric state of the purified mini-COL4A345 complex sample were assessed by sedimentation velocity analytical ultracentrifugation (SV-AUC) on a Beckman Coulter Optima analytical ultracentrifuge. The sample was prepared in phosphate buffered saline (PBS) and loaded into a two-sector Epon centrepiece fitted with quartz windows, with matched buffer in the reference sector. Cells were placed in an An-50 Ti rotor and sedimentation was carried out at 163,000 g at 20 °C, with radial absorbance scans recorded at 230 nm at 60 s intervals throughout the run. The protein partial specific volume and the buffer density and viscosity were calculated in SEDNTERP [16, 17]. Data were analysed in SedFit [18] using a continuous sedimentation coefficient [c(s)] distribution model, with sedimentation coefficients corrected to standard conditions (s₂₀,w). Molecular masses were determined within SedFit via the Svedberg equation using the fitted frictional ratio (f/f₀).

### Mass photometry

The molecular weights of the mini-COL4A345 complex were determined by mass photometry using a Refeyn TwoMP mass photometer. Samples were diluted to approximately 33 nM in PBS immediately prior to measurement. After focussing on buffer alone, the sample was applied and the binding ("landing") of individual molecules onto the glass coverslip was recorded as interferometric scattering contrast over a 60 s acquisition. The contrast of each single-molecule binding event, which scales linearly with molecular mass, was measured; the resulting contrast values were binned into a histogram and fitted with Gaussian functions to resolve the individual mass populations. Contrast was converted to molecular mass using a three-point calibration from protein standards of known molecular weight bovine serum albumin monomer (66 kDa) and dimer (132 kDa), and alcohol dehydrogenase tetramer (147 kDa) acquired under identical conditions. Data were collected and analysed using Refeyn AcquireMP and DiscoverMP software, respectively.

### AlphaFold structure prediction

Protein structure prediction was performed using AlphaFold2, a deep learning-based protein structure prediction algorithm [19]. The amino acid sequence of the recombinant human mini-COL4A345 protomer was used as input (**Supplementary Figure 1**), and structural models were generated using the AlphaFold Protein Structure Database. Structural visualisation and analysis were performed using PyMOL.

### Mouse monoclonal antibodies

Monoclonal antibodies against mini-COL4A345 were generated by a commercial service provider (ProteoGenix, France) using antigen supplied by the authors. Mouse immunisations and spleen collection for monoclonal antibody generation were performed by ProteoGenix (France) in accordance with European Directive 2010/63/EU concerning the protection of animals used for scientific purposes. The procedures were conducted under appropriate institutional ethical approval and authorisation. Mice were immunised with the mini- COL4A345 antigen and splenocytes were fused with a myeloma cell line to generate hybridomas. Hybrid cells were selected at the polyclonal stage and expanded for ELISA screening against the purified antigen. The best-performing hybridomas were subsequently screened and candidate clones were validated in-house.

### Hybridoma screening by biolayer interferometry and immunoassays

Hybridoma culture supernatants were screened for binding to recombinant COL4A345 antigen using biolayer interferometry (BLI) on an Octet Red96e instrument at a set temperature of 30 °C. Hybridoma supernatants were diluted 1:2 in a running buffer (RB) of PBS pH 7.4 containing 0.05% Tween-20 (PBS-T) and captured onto protein A biosensors. Protein A sensors without captured antibody were used as reference sensors. Following antibody capture and baseline equilibration in RB, biosensors were immersed in either 100 nM recombinant mini-COL4A345 antigen or PBS-T alone as a negative control. Association and dissociation responses were recorded, and dissociation rate constants (*k*_d_, s^−1^) were determined using Octet Data Analysis HT software. Antibodies showing binding to COL4A345 with minimal non-specific binding were selected for further characterisation.

Immunoassays were used to assess hybridomas reactivity with COL4 proteins under native dot blotting or denaturing western blotting conditions. For dot blotting, 0.5 µg of purified human protein (mini-COL4A345, -COL4A3 or -COL4A5) was blotted onto nitrocellulose membrane under non-reducing conditions using a Minifold I dot-blot system (Fisher UK, BRD-600-025S). For western blotting, 0.5 µg of purified human protein was loaded under reducing conditions onto a 4–12% NuPAGE gel (Thermo Fisher, NP0321) and transferred for 70 min at 30 V onto nitrocellulose membrane using the Invitrogen XCell SureLock Mini-Cell and XCell II Blot Module system (Thermo Fisher, 10572913). One membrane was probed per hybridoma. Membranes were blocked with Intercept blocking buffer (LI-COR, 927- 70001) for 1 h at room temperature, incubated with hybridoma supernatants diluted 1:4 in PBS-T overnight at 4 °C, washed three times in PBS-T, and incubated with anti-mouse IgG DyLight 800 secondary antibody (1:10,000; Cell Signaling Technology, 5257S) for 1 h at room temperature. Membranes were washed three times in PBS-T and imaged using an Odyssey CLx Imager (LI-COR).

### Binding analysis of purified antibodies

Surface plasmon resonance (SPR) analyses were carried out on a Biacore™ 1K+ (Cytiva) using a Series S CM5 sensor chip with an assay running buffer of HBS with 0.05% Tween20, pH7.4. Reference (flow cell 1) and active flow cells were activated by injecting a 1:1 mixture of 0.2 M 1-ethyl-3-(3-dimethylaminopropyl)-carbodiimide (EDC) and 0.1 M N- hydroxysuccinimide (NHS) at 10 µL/min for 420 seconds. Mini-COL4A345 protein was immobilised onto flow cell 2 at 10 µg/ml in 10 mM sodium acetate pH 4.0 at 10 µL/min to approximately 200 response units (RU). COL4A3 and COL4A5 were immobilised onto flow cells 3 and 4, respectively, at 1 µg/ml in 10 mM sodium acetate pH 5.5 at 10 µL/min to approximately 100 RU. Reference and active flow cells were blocked using 1 M ethanolamine, pH 8 at a flow rate of 10 µL/min for 420 seconds. For kinetic analysis, antibodies were flowed over active and reference flow cells at 30 µL/min in assay running buffer at concentrations of 3.7 - 300 nM for Ms#72 and 12.3 - 1000 nM for Ms#5 and Ms#44, using a three-fold dilution series. Single-cycle kinetics were performed with a contact time of 120 seconds and a 900 second dissociation time. Analysis temperature was set at 37 °C. Surfaces were regenerated between antibody samples with 2x 5 second injections of 0.1 M Glycine, pH 2.0 at a flow rate of 30 µL/min. Reference subtracted data sets were analysed in the Biacore™ Insight Evaluation software using the bivalent analyte model.

### Mouse models

*Col4a5* knockout mice (*Col4a5^−/Y^*), an X-linked Alport syndrome model, were obtained from the International Mouse Phenotyping Consortium [20] and bred on a C57BL/6N background [21]. Mice were euthanised for tissue collection in compliance with UK Home Office Animals (Scientific Procedures) Act 1986. Mouse kidney tissues from *Col4a3* knockout mice [22] were kindly provided by collaborators at Washington University School of Medicine. These animals were euthanised for tissue collection in accordance with the National Institutes of Health Guide for the Care and Use of Laboratory Animals, under protocols approved by the Washington University Institutional Animal Care and Use Committee (IACUC). Mouse kidney tissues from *Col4a6* knockout mice [23] were kindly provided by collaborators at Okayama University. Euthanasia for tissue collection was conducted in accordance with protocols approved by the institutional Animal Care and Use Committee at Okayama University (approval no. OKU-2024134).

### Human kidney tissue procurement

Human kidney tissue was obtained through the MFT Biobank, which operates under ethical approval from the NRES Committee North West – Haydock (REC 19/NW/0644). Healthy control kidney tissue was obtained from the non-diseased portion of nephrectomy samples.

### Kidney immunofluorescence

Formalin-fixed paraffin-embedded (FFPE) mouse kidneys (wild-type C57BL/6N, *Col4a5* knockout, *Col4a3* knockout and *Col4a6* knockout) were sectioned at 3 µm using a Leica RM2255 microtome (n = 1 biological sample per condition). Tissue sections were dewaxed and rehydrated. Antigen retrieval was performed using DAKO citrate buffer, pH 6 (Agilent, S236984-2), heated in a benchtop steamer to 95 °C for 30 min, followed by cooling at room temperature for 30 min. Flash-frozen mouse kidneys were embedded in OCT, sectioned at 5 µm using a Leica CM1950 cryostat and fixed in acetone at −20 °C for 5 min. Sections were washed in PBS and blocked with avidin/biotin solution (Vector Laboratories, SP-2001), followed by M.O.M. Mouse IgG Blocking Reagent (Mouse on Mouse Immunodetection Kit, Fluorescein; Vector Laboratories, FMK-2201) for 1 h at room temperature. Sections were washed in PBS and prepared for antibody incubation. Paraffin-embedded human kidneys were cut to 3 μm thickness using a Leica RM2255 microtome. Heat-induced epitope retrieval was performed by incubating tissue sections in 10 mM tri-sodium citrate buffer pH 2, 0.05% Tween-20 for 20 min at 95 °C. Sections were washed with water before incubating in 0.1M Glycine, 6M urea pH3.2 buffer for 30 mins at room temperature. Tissue was washed with PBS then blocked with DAKO blocking buffer (Agilent, X0909) for 1 hour. Sections were washed in PBS and prepared for antibody incubation. The primary antibodies (purified antibodies clones #5, #44 and #72) were diluted at 10 µg/ml in DAKO antibody diluent with background-reducing buffer (Agilent, #S302281-2) and incubated overnight at 4°C in a humidified chamber. For co-staining, rat collagen IV chain-specific antibodies (α1–α6) antibodies (1:200; Chondrex, COL4A1 H11 #7070, COL4A2 H22 #7071, COL4A3 H31 #7076, COL4A4 H43 #7073, COL4A5 H52 #7077, COL4A6 H63 #7074) were added. The secondary antibodies anti-mouse Alexa Fluor 594 (1:400; Invitrogen, #A-11005) and anti-rat Alexa Fluor 488 (1:400; Invitrogen, #A48262TR) were incubated for 45 min at room temperature in a humidified chamber. After washing with PBS, slides were dried and mounted with ProLong Gold Antifade Mountant (Thermo Fisher, P36930). Images were collected using a Zeiss LSM 880 upright Airyscan confocal microscope. Immunofluorescence staining was performed twice for each sample, with consistent staining patterns observed between experiments.

### Cochlear immunofluorescence

To assess protein localisation in the cochlea, wild-type C57BL/6N and *Col4a5* knockout mice at postnatal day 21 were culled by cervical dislocation (n = 1 biological sample per condition). Inner ears were removed and fixed by perfusion through the round and oval windows with 4% paraformaldehyde, followed by 1 h submersion fixation on ice. Whole tissues were decalcified for 24–72 h at 4 °C in 4% EDTA, cryoprotected with 30% sucrose, embedded in OCT and sectioned at 12 µm using a cryostat. Sections were post-fixed on slides in 4% paraformaldehyde for 5 min at room temperature, permeabilised with PBS containing 0.2% Triton X-100 and blocked in 5% goat serum. Endogenous mouse IgG binding was blocked using ReadyProbes Mouse-on-Mouse IgG Blocking Solution (Invitrogen, R37621) before incubation with Ms#72 at 5 µg/ml in DAKO antibody diluent with background-reducing buffer overnight at 4 °C. Goat anti-mouse IgG (H+L) Alexa Fluor 488 secondary antibody (1:500; Thermo Fisher Scientific, A-11029) was applied for 1 h at room temperature, followed by a 5 min incubation with DAPI (300 nM; Thermo Fisher Scientific, 62248). Slides were mounted using ProLong Glass Antifade Mountant (Invitrogen, P36980) and imaged using a Zeiss LSM 880 confocal microscope or Zeiss Axio Imager.M2 fluorescence microscope. Images were processed using Zeiss Zen microscopy software and Fiji. Immunofluorescence staining was performed twice for each sample, with consistent staining patterns observed between experiments.

### Eye immunofluorescence

18-week-old wild-type C57BL/6N mouse eyes were retrieved, embedded in OCT and sectioned on a Leica CM1950 cryostat into 10 μm slices (n = 1 biological sample per condition). Sections were immersed in ice-cold acetone for 10 min followed by 10 min wash in PBS. Sections were blocked in PBS containing 0.05% Tween20 and 0.2% fish gelatin for 30 min. Sections were incubated overnight at 4°C with Ms#72 conjugated with FlexAble CoraLite® Plus 594 fluorophore (ProteinTech, KFA049) and rabbit anti-panLaminin (1:400; Abcam, ab11575). Briefly, 0.5 µg of primary Ms#72 was combined with 1 µL of FlexLinker, and FlexBuffer was added to a final volume of 8 µL. The mixture was gently mixed and incubated for 5 min at room temperature protected from light. Subsequently, 2 µL of FlexQuencher was added, followed by gentle mixing and a further 5 min incubation at room temperature in the dark. The labelled antibody was then diluted 1:5 in DAKO antibody diluent and added to the samples. After incubation, sections were washed 3 times for 5 min in PBS, followed by incubation with anti-rabbit Alexa Fluor 488 secondary antibodies (1:400; Invitrogen, A32731TR) for 40 min at room temperature. Slides were incubated with DAPI before being mounted using EverBrite non-curing mounting medium (Biotium, #23001) and imaged using the LSM 880 confocal. Immunofluorescence staining was performed twice for each sample, with consistent staining patterns observed between experiments.

### Immunoprecipitation of kidney tissue using Ms#72

Kidneys from wild-type and *Col4a5* knockout mice were harvested, rinsed in cold TBS and minced finely with scissors. Kidneys were homogenised with a pestle in cold TBS containing protease inhibitors (Roche, 04693159001) and centrifuged at 4000 × *g* for 5 min at 4 °C. The pellet was washed three times with TBS. To examine NC1 hexamer crosslinks, the pellet was resuspended in digestion buffer containing 50 mM Tris-HCl, pH 7.4, 150 mM NaCl, 5 mM CaCl_2_, 10 µg/mL DNase I, 0.1% Triton X-100, protease inhibitors, 2 mg/mL collagenase XI (Sigma-Aldrich, C9407) and 1 mg/mL Liberase TL (Roche, 05401020001). Samples were incubated at 37 °C overnight with rotation. The reaction was stopped by adding 10 mM EDTA and centrifuged at 6000 × *g* for 15 min. The supernatant containing soluble NC1 complexes was collected and incubated with 40 µL Dynabeads Protein G (Invitrogen, 10003D) for 2 h at 4 °C with rotation to pre-clear the sample. Magnetic beads were removed, and the supernatant was incubated overnight at 4 °C with 1 µg Ms#72 antibody. The next day, 40 µL Dynabeads Protein G was added and samples were incubated for 2 h at 4 °C with rotation. Immunocomplexes were eluted with 50 mM glycine, pH 2.8, and analysed by SDS-PAGE using a 4–12% gradient gel under non-reducing conditions. Proteins were transferred onto nitrocellulose membranes and incubated overnight at 4 °C with rat anti- COL4A3 NC1 antibody (1:1000; clone H31, Chondrex, 7076), rat anti-COL4A4 NC1 antibody (1:1000; clone H43, Chondrex, 7073), rabbit anti-COL4A5 NC1 antibody (1:1000; BiCell, 05005), or rat anti-COL4A6 NC1 antibody (1:1000; clone H63, Chondrex, 7074). Membranes were incubated with goat anti-rabbit IgG DyLight 800 (1:10,000; Cell Signaling Technology, 5151) and goat anti-rat Alexa Fluor 680 (1:10,000; Invitrogen, A-21096). Bands were detected using a ChemiDoc imaging system.

## Acknowledgments

We gratefully acknowledge Tomoko Yonezawa and Toshitaka Oohashi from Okayama University (Japan) for providing the *Col4a6* knockout mouse tissue and Jeffrey Miner of Washington University in St Louis (USA) for providing the *Col4a3* knockout mouse tissues.

## Data availability

All data described in this manuscript and supporting information are contained within the manuscript or are available from the corresponding author upon reasonable request.

## Funding and additional information

This work was supported by Four Points Innovation, a portfolio company of investment funds managed by Deerfield Management Company, L.P. D.J.J is supported by Kidney Research UK (ARP/001/20240902) and a Biotechnology and Biological Sciences Research Council grant (BB/R000549/1). R.L. is funded by the Wellcome Trust (Ref: 226804/Z/22/Z, Ref: 227417/Z/23/Z, Ref: 301803/Z/23/Z); Kidney Research UK and The Stoneygate Trust (Alport Research Hub), the National Institute for Health Research (NIHR-Manchester BRC), and Visterra Therapeutics. She has consultancy agreements with Calliditas Therapeutics, River 3 Renal and Travere Therapeutics.

---

**Supplementary Figure 1. Mini-COL4A345 antigen design and validation.** Annotated plasmid maps for mini-COL4A3, mini-COL4A4 and mini-COL4A5 expression constructs.

**Supplementary Figure 2. Mass photometry assessment of mini-COL4A345 oligomerisation.** Mass photometry was used to assess the oligomeric state of purified mini- COL4A345 in solution.

**Supplementary Figure 3. Hybridoma screening and monoclonal antibody selection.** A) ELISA screening of hybridomas against purified mini-COL4A345 antigen. Positive hybridomas are highlighted in yellow. Red highlights denote hybridomas identified by BLI screening. Solid lines indicate “high” binders and dashed lines indicate “low” binders based on binding response and signal above reference-subtracted baseline. B) Hybridoma selection workflow showing the screening process and outcome from BLI, denaturing Western blotting, native dot blotting and immunofluorescence on human kidney tissue, resulting in the selection of six hybridomas. Scale bar, 100 µm.

**Supplementary Figure 4. Wild-type mouse kidney tissue labelled with the three selected hybridomas.** Mouse kidney FFPE sections (A) and cryosections (B) labeled with hybridomas Ms#5, Ms#44 and Ms#72. Ms#72 showed the strongest signal in the glomerular basement membrane with the lowest background. Scale bar, 50µm.

**Supplementary Figure 5. Co-localisation analysis of Ms#72 in human kidney tissue.** FFPE kidney sections from a healthy control were co-stained with Ms#72 and antibodies against COL4A1 or COL4A2. Scale bar, 50µm.

