## Supplementary figures for "Monoclonal antibodies for detecting collagen IV α3α4α5 in basement membranes"

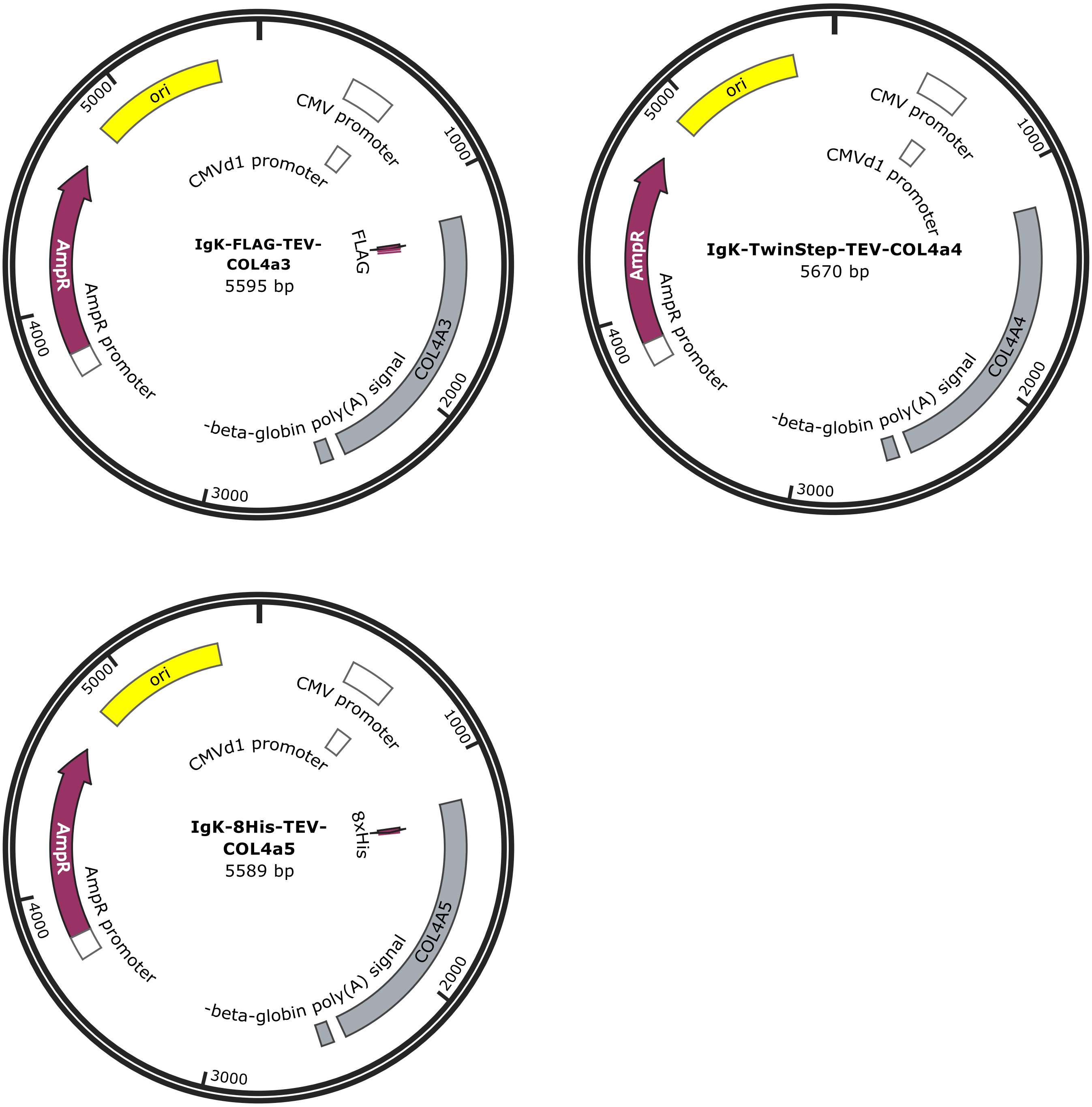


**Supplementary Figure 1. Mini-COL4A345 antigen design and validation.**
Annotated plasmid maps for mini-COL4A3, mini-COL4A4 and mini-COL4A5 expression constructs.

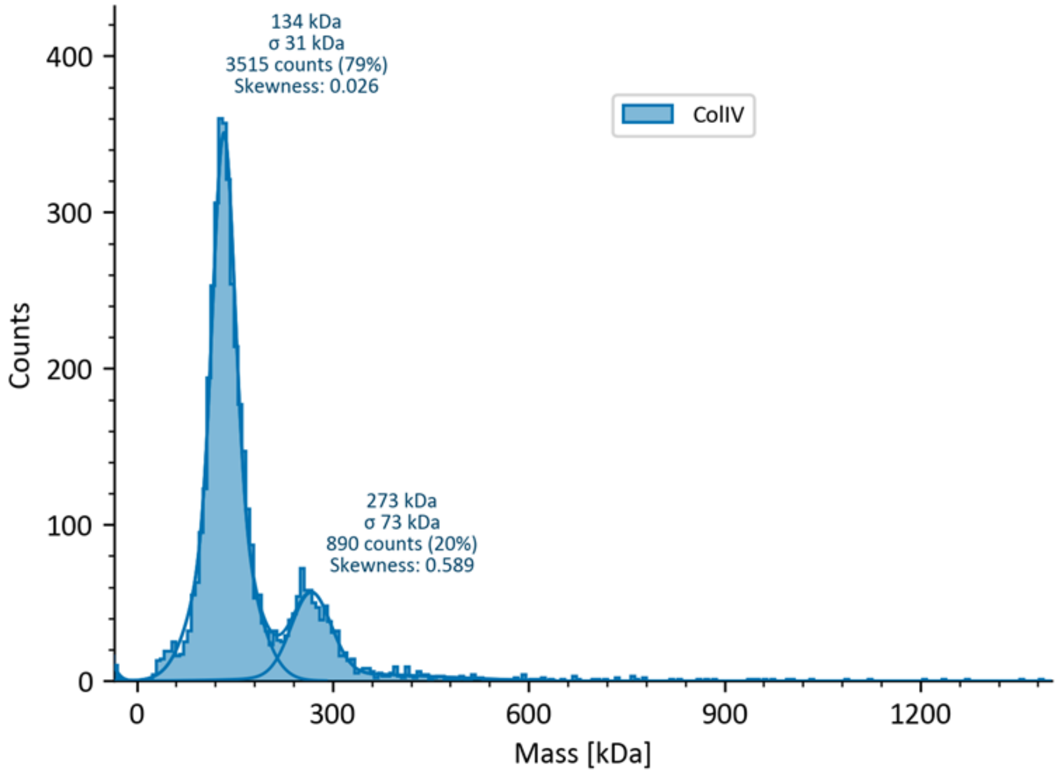


**Supplementary Figure 2. Mass photometry assessment of mini-COL4A345 oligomerisation.** Mass photometry was used to assess the oligomeric state of purified mini-COL4A345 in solution.

**A**

| Confirmation screening after fusion for Mouse #3 #4 #5 | | | | | | | | | | | | |
|---|---|---|---|---|---|---|---|---|---|---|---|---|
| Clone ID | 1 | 2 | 3 | 4 | 5 | 6 | 7 | 8 | 9 | 10 | 11 | 12 |
| Col IV heteroprotomer | 2,75 | 1,68 | 2,74 | 2,77 | 3,24 | 0,98 | 1,90 | 1,11 | 2,37 | 2,49 | 1,52 | 2,28 |
| Clone ID | 13 | 14 | 15 | 16 | 17 | 18 | 19 | 20 | 21 | 22 | 23 | 24 |
| Col IV heteroprotomer | 1,40 | 1,04 | 1,63 | 1,22 | 1,38 | 1,47 | 1,47 | 1,74 | 2,03 | 2,62 | 2,05 | 0,03 |
| Clone ID | 25 | 26 | 27 | 28 | 29 | 30 | 31 | 32 | 33 | 34 | 35 | 36 |
| Col IV heteroprotomer | 1,62 | 0,96 | 1,19 | 1,07 | 2,89 | 1,74 | 1,17 | 1,31 | 2,53 | 1,85 | 2,47 | 1,59 |
| Clone ID | 37 | 38 | 39 | 40 | 41 | 42 | 43 | 44 | 45 | 46 | 47 | 48 |
| Col IV heteroprotomer | 2,44 | 2,13 | 2,45 | 2,82 | 1,64 | 1,52 | 2,39 | 2,65 | 1,61 | 1,41 | 2,54 | 2,27 |
| Clone ID | 49 | 50 | 51 | 52 | 53 | 54 | 55 | 56 | 57 | 58 | 59 | 60 |
| Col IV heteroprotomer | 1,70 | 1,81 | 2,11 | 2,72 | 2,63 | 1,72 | 1,33 | 1,37 | 1,10 | 1,87 | 1,21 | 1,71 |
| Clone ID | 61 | 62 | 63 | 64 | 65 | 66 | 67 | 68 | 69 | 70 | 71 | 72 |
| Col IV heteroprotomer | 0,63 | 2,71 | 1,20 | 2,76 | 2,45 | 2,12 | 0,78 | 2,76 | 1,30 | 1,97 | 2,63 | 2,15 |
| Clone ID | 73 | 74 | 75 | 76 | 77 | 78 | 79 | 80 | 81 | 82 | 83 | 84 |
| Col IV heteroprotomer | 3,18 | 1,58 | 2,01 | 1,79 | 2,06 | 1,72 | 2,23 | 2,16 | 1,28 | 1,10 | 1,83 | 0,74 |
| Clone ID | 85 | 86 | 87 | 88 | 89 | 90 | 91 | 92 | 93 | 94 | negative | positive |
| Col IV heteroprotomer | 3,23 | 0,71 | 0,72 | 0,78 | 1,37 | 1,01 | 1,34 | 2,24 | 1,17 | 0,61 | 0,03 | 3,05 |

**B**

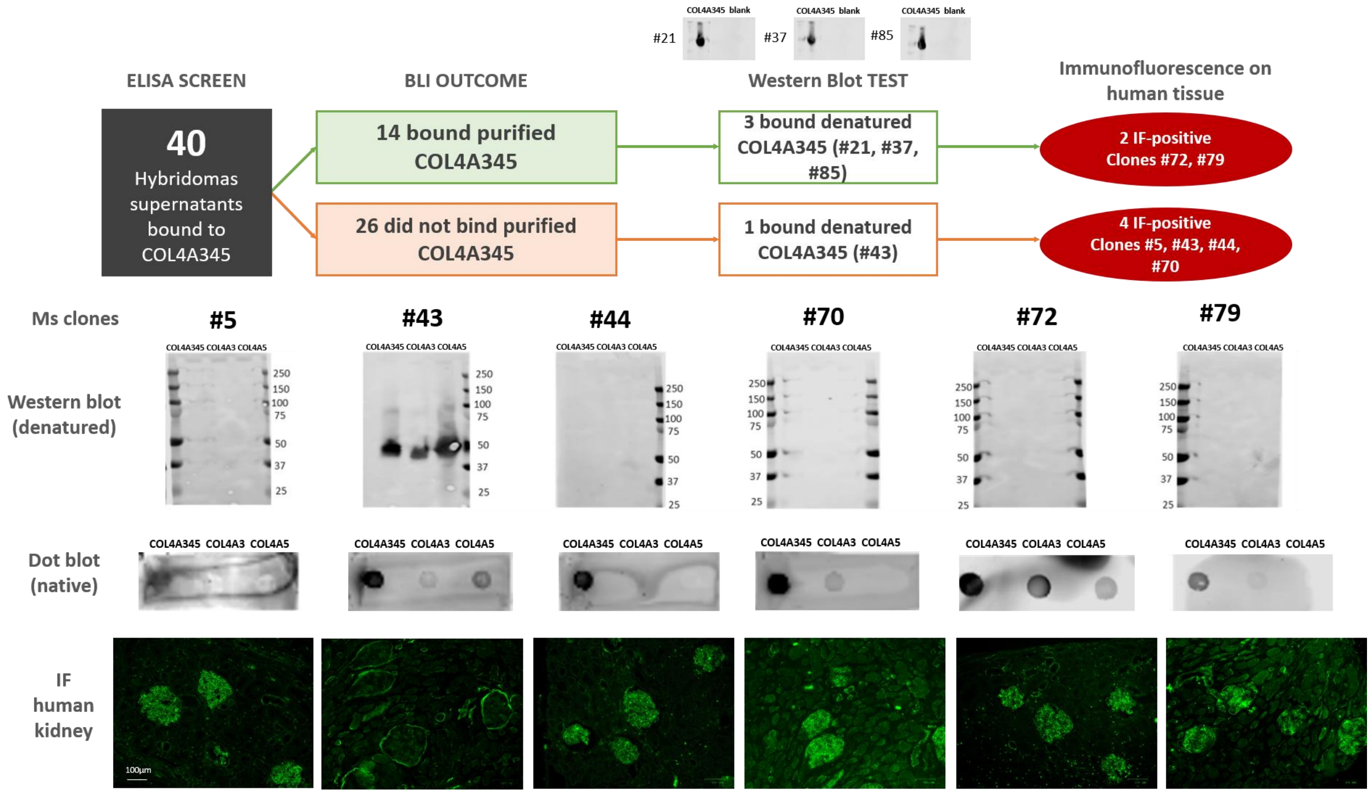


**Supplementary Figure 3. Hybridoma screening and monoclonal antibody selection.**
A) ELISA screening of hybridomas against purified mini-COL4A345 antigen. Positive hybridomas are highlighted in yellow. Red highlights denote hybridomas identified by BLI screening. Solid lines indicate “high” binders and dashed lines indicate “low” binders based on binding response and signal above reference-subtracted baseline. B) Hybridoma selection workflow showing the screening process and outcome from BLI, denaturing Western blotting, native dot blotting and immunofluorescence on human kidney tissue, resulting in the selection of six hybridomas. Scale bar, 100 µm.

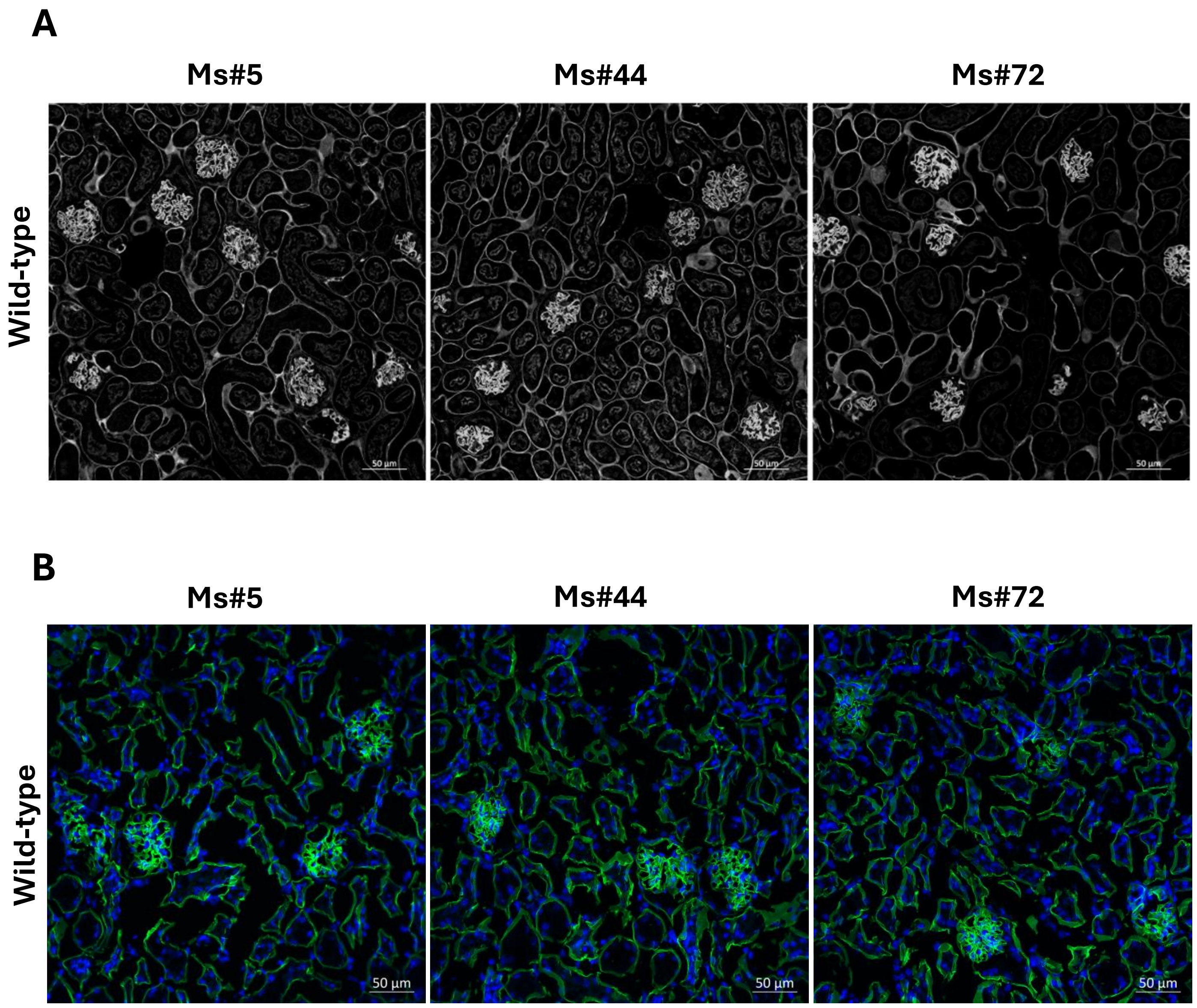


**Supplementary Figure 4. Wild-type mouse kidney tissue labelled with the three selected hybridomas.** Mouse kidney FFPE sections (A) and cryosections (B) labeled with hybridomas Ms#5, Ms#44 and Ms#72. Ms#72 showed the strongest signal in the glomerular basement membrane with the lowest background. Scale bar, 50µm.

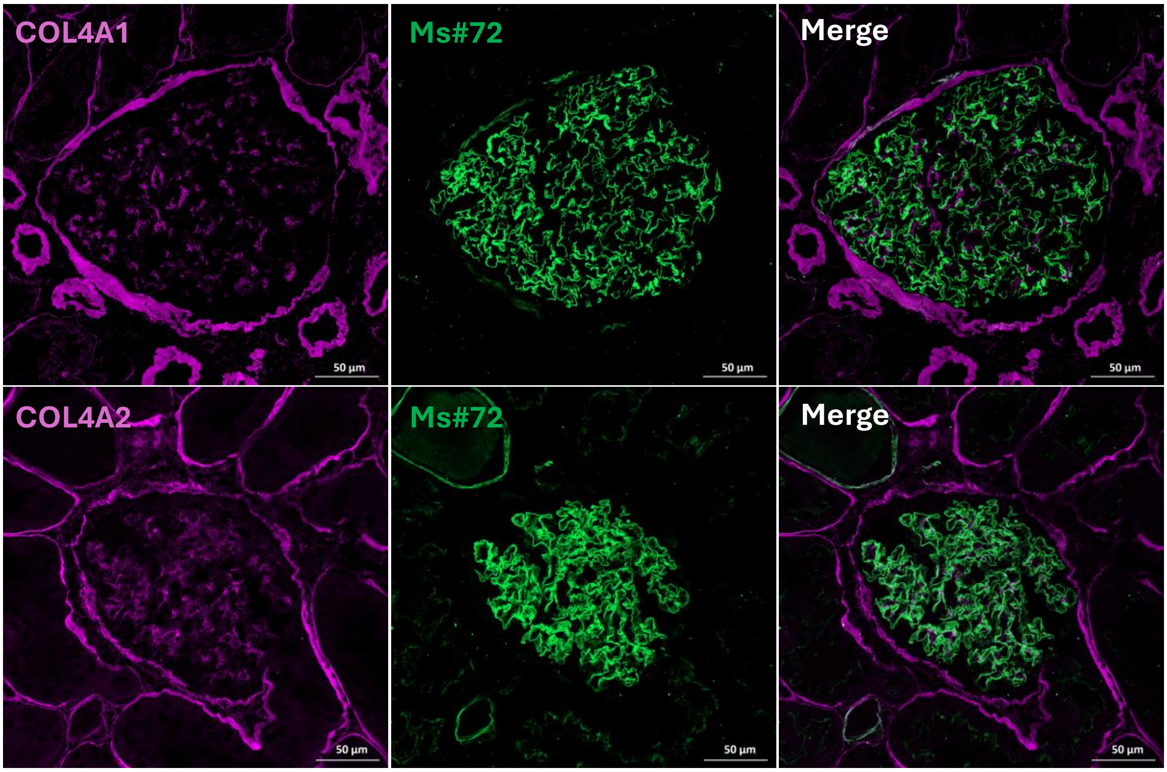


**Supplementary Figure 5. Co-localisation analysis of Ms#72 in human kidney tissue.** FFPE kidney sections from a healthy control were co-stained with Ms#72 and antibodies against COL4A1 or COL4A2. Scale bar, 50μm.
